# Fibrotic pulmonary dust foci is an advanced pneumoconiosis lesion in rats induced by titanium dioxide nanoparticles in a 2-year inhalation study

**DOI:** 10.1101/2024.12.19.629539

**Authors:** Shotaro Yamano, Yumi Umeda

## Abstract

**Background:** We have previously reported that inhalation exposure to titanium dioxide nanoparticles (TiO_2_ NPs) for 13 weeks causes early pneumoconiosis lesions in the alveolar region of F344 rats. We defined these characteristic lesions as pulmonary dust foci (PDF). In this report, we re-evaluate and detail the histopathological data regarding particle-induced pneumoconiosis lesions, including progressive lesions of the early PDF lesions, that developed in F344 rats exposed TiO_2_ NPs by whole body inhalation over a period of two years.

**Methods:** Male and female F344 rats were exposed to 0.5, 2, and 8 mg/m^3^ anatase type TiO_2_ NPs for 6 hours/day, 5 days/week for 104 weeks using a whole-body inhalation exposure system. After the final exposure, the rats were euthanized. In the present study, the collected lungs were re-evaluated macroscopically and histopathologically.

**Results:** Rats exposed to TiOL NPs developed macroscopic white lesions, primarily in the subpleural and hilar regions of the lung, which increased in size and number with exposure concentration. Histologically, two lesion types were identified: (1) Fibrotic Pulmonary Dust Foci (fPDF), characterized by collagen deposition, inflammatory infiltration, and disrupted alveolar epithelial differentiation, and (2) Dust Macules (DM), characterized by macrophage accumulation without significant fibrosis or inflammation. Immunohistochemical analysis revealed abnormal alveolar epithelial differentiation and reduced capillary density within fPDFs. Importantly, no histological connection was found between the pneumoconiosis lesions and the observed lung tumors, which resembled spontaneous, age-related neoplasms.

**Conclusions:** Chronic inhalation of TiOL NPs induces advanced pneumoconiosis characterized by fPDF and DM, with distinct pathological features. However, these lesions were not directly linked to lung tumor development. Therefore, in this study PDF lesions developed into fPDF lesions but did not lead to tumorigenesis. This study provides critical insights into the long-term pulmonary effects of TiOL NP exposure and the progression of pneumoconiosis lesions in the rats.

## Background

Pneumoconiosis is well-understood pathologically as an airspace and/or interstitial lung disease caused by particle-laden macrophages; pneumoconiosis is chronic, progressive and still has no fundamental treatment [1–5]. Furthermore, the progression of pneumoconiosis is well known to increase the risk of lung cancer [6,7]. It is known from previous epidemiology and case reports of workers that pneumoconiosis can be caused by inhalation of various materials, including asbestos [8], silica [9], mixed dust [10], hard metals [11,12], aluminum [13], beryllium [14], indium [15–17], talcum [18], and also titanium dioxide (TiO_2_)/titania particles including nanoparticles (NPs) and titanium grindings [19–30]. However, most of the research conducted to date on laboratory animals using nano/micro particles, including workplace environment particles and nanomaterials, has focused on acute toxicity and carcinogenicity, and there have been few pathological findings based on the development of pneumoconiosis.

In recent years, we have conducted whole-body inhalation exposure studies using rodents exposed to indium tin oxide particles[31], multi-walled carbon nanotubes[32,33], TiO_2_ NPs[34,35], and cross-linked water-soluble acrylic acid polymers[36,37], and have carried out detailed histopathological analysis of lesions induced in the rodent lung. In accordance with the Organization for the Economic Co-operation and Development Guideline for Testing of Chemicals 413 (OECD TG 413)[38], we conducted a 13-week inhalation toxicity study using 100 male and female F344 rats, and found that inhalation of anatase-type TiO_2_ NPs induced early pneumoconiosis lesions in the lungs, which we defined as pulmonary dust foci (PDF)[35]. Histopathological examination showed that the PDF consisted of aggregations of macrophages that had fully ingested particles in the alveolar space and hyperplasia of alveolar type L cells (AT2) around the macrophages, but no increase in collagen fibers in the alveolar septa was observed. A two-year inhalation toxicity study of the same anatase-type TiO_2_ NPs was then conducted in accordance with the OECD TG 451[39], which focused on carcinogenicity[40]. However, progression of pneumoconiosis lesions, including the progression of the disease state from PDF, has not been fully investigated.

Therefore, the aim of this study was to re-evaluate the development and progression of pneumoconiosis lesions by conducting a more detailed macroscopic and histopathological evaluation of lung tissue samples obtained from a two-year carcinogenicity study.

## Results

### Representative macroscopic findings of TiO_2_ NPs-exposed rat lung

The overall experimental protocol for the re-evaluation of the development and progression of pneumoconiosis lesions is shown in Fig. 1A. Representative macroscopic findings are shown in Fig. 1B-M. Numerous white clusters were observed in the lungs of male and female rats exposed to all three concentrations of TiO_2_ NPs (0.5, 2, 8 mg/m^3^), and both number and size were increased in a concentration-dependent manner (Fig. 1B-G). Notably, females exhibited a greater number of white clusters than males at each exposure concentration. The white clusters were primarily located in the periphery on the dorsal (rib) side of each lung lobe (Fig. 1H and 1I). Further magnification identified two distinctive types of changes in the white clusters, clusters with numerous white-gray spots (1-2 mm in size, Figs. 1J, 1L, 1M) and clusters consisting of small pure white spots (less than 500 μm in size, Figs. 1J and 1K).

**Figure 1:**
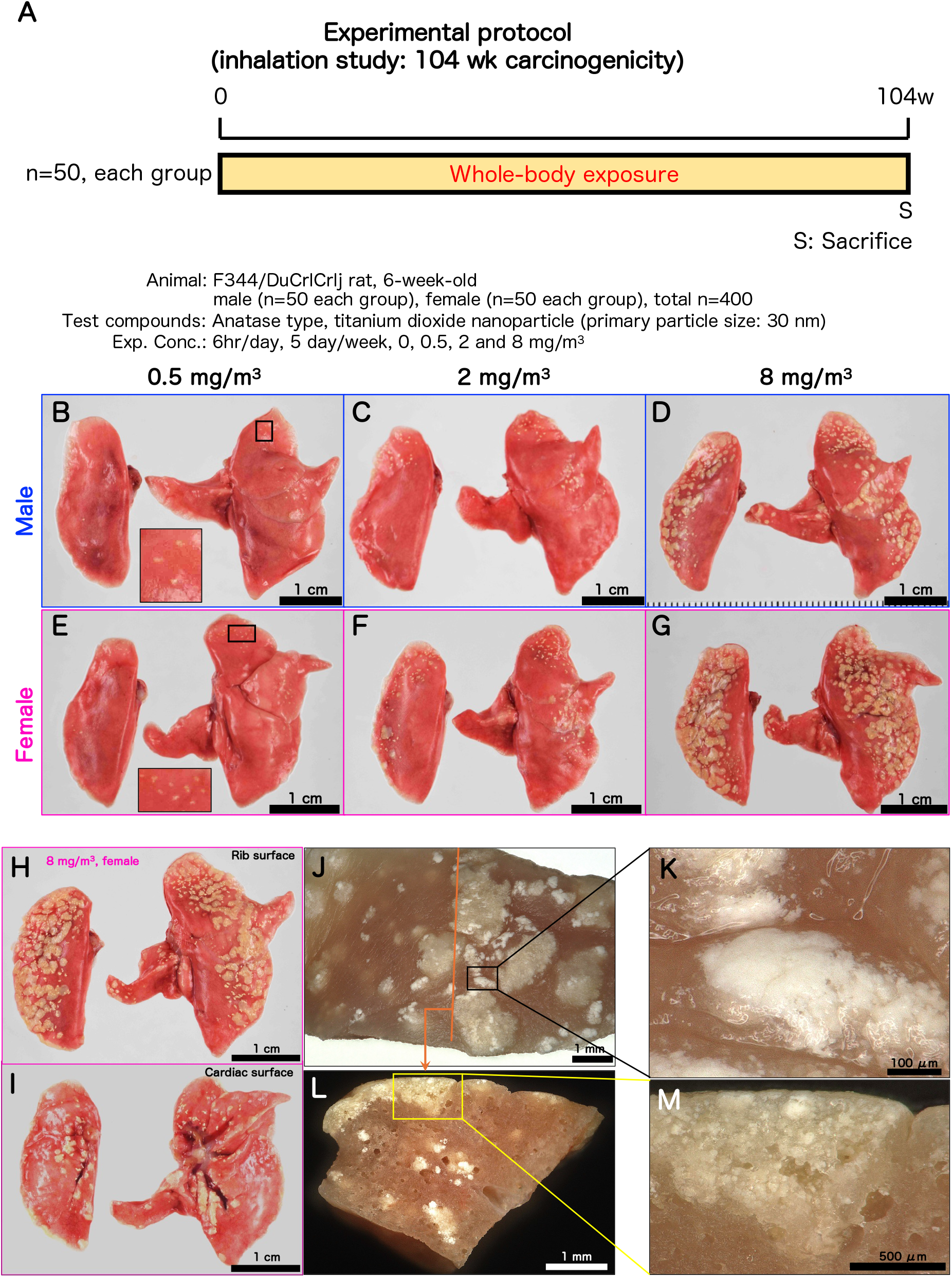
Experimental protocol and representative macroscopic photographs of the lungs of F344 rats after 2-year whole body inhalation exposure to TiO_2_ NP. A total of 400 F344 rats were used in the experimental protocol (A) for the two-year whole-body inhalation exposure experiment. Representative macroscopic photographs of the lungs immediately after the final dissection are shown for male and female rats in the 0.5 mg/m^3^ group (B, E), 2 mg/m^3^ group (C, F), and 8 mg/m^3^ group (D, G). Bar: 1 cm. For female rats in the 8 mg/m^3^ exposure group, macroscopic photographs of the rib surface (H) and cardiac surface (I) of the same animal showed the characteristic localization of white clusters. Bar: 1cm. Enlarged photographs (J, bar: 1 mm) and cross-sectional photographs (L, bar: 1 mm) of white clusters show that they can be distinguished into two characteristic foci: white-gray spots (M, bar: 500 μm) and small pure white spots (K, bar: 100 μm). Fig.1I shows a reversed image. High resolution macroscopic images were captured by VHX-8000 digital microscope (Keyence, Osaka, Japan).

Additionally, fusions of white-gray spots and small pure white spots were observed in dense areas (approximately 3 mm in size) (Figs. 1L and 1M). Furthermore, in the areas where the four lobes of the right lung overlap, a decrease in all the spots was observed.

These results suggest that prolonged exposure to inhaled TiO_2_ NPs for 2 years leads to the development of a substantial number of clustered lesions in characteristic localizations, in a concentration-dependent manner, in the lungs of rats.

### Re-evaluation of histopathological characteristics of TiO_2_ NPs-exposed rat lung

Formalin fixed lungs were cut as illustrated in Fig. S1, and embedded in paraffin. Serial sections were then prepared for histological examination. Lung lesions were predominantly subpleural and multifocal, consistent with the areas where white clusters were visible on gross examination (Fig. 2A). Lesions within and along the airways in the bronchial and bronchiolar regions were rare, with the main pathological changes occurring in the alveolar regions.

**Figure 2:**
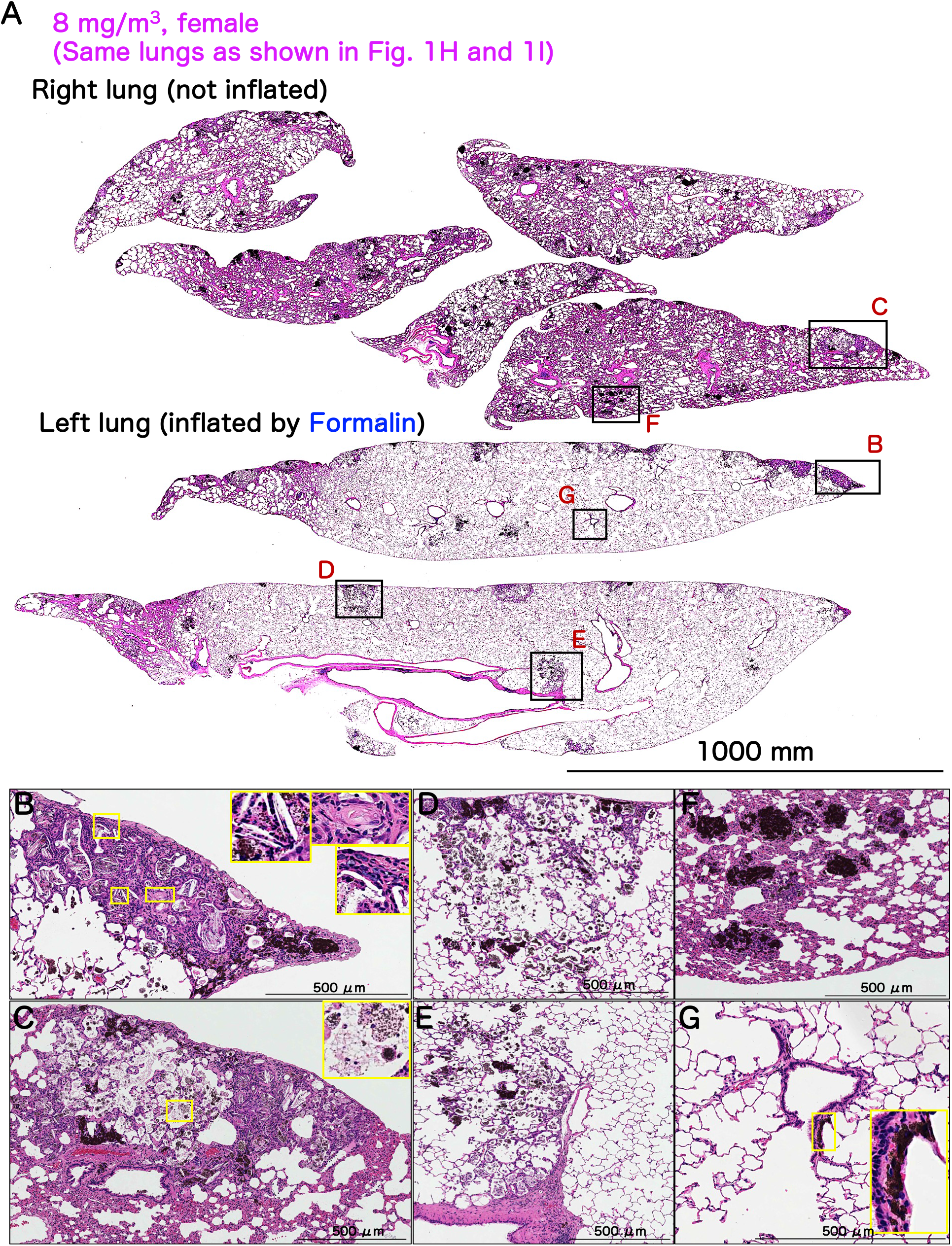
Representative microscopic photographs of female rat lungs after inhalation exposure to TiO_2_ NP (8 mg/m^3^). Loupe image (A, bar: 1000 mm) and magnified image (B-G, bar: 500 μm) of H&E-stained lung sections of the lungs shown in Figs. 1H and I. White-gray spots macroscopically detected were observed as nodular lesions with relatively clear boundaries (B and right side of C). Severe lesions tended to be scattered in the apex of the lung. The lesions were composed of cholesterol clefts, collagen bundles, and mast cells (enlarged in B). Lesions with mild changes were scattered below the apex of the lung (right side of C) in the subpleural area (D) and the hilum (E). The alveolar structure was well preserved, and the macrophages in the alveolar airspace were mostly destroyed (enlarged in C). The small pure white spots macroscopically detected were observed as clumps of black pigment (F) and were foci of aggregations of particle-phagocytosing macrophages. In the interstitium of the terminal bronchioles, also known as the broncho-vascular bundle, the deposition of particle-phagocytosing macrophages was mild (G).

The lesions observed macroscopically as white-gray spots were caused by a variety of lesions with varying severity. Around the apical lung area, lesions were characterized by the destruction/distortion of existing alveolar structures with remodeling of the tissue architecture, forming nodular lesions with cholesterol granulomas with multinucleated giant cells, collagen fiber bundles, and mast cells, often accompanied by hyperplasia of the epithelium (Figs. 2B, 2C-right region). In contrast, lesions in the subpleural areas other than the apex of the lung (Fig. 2D) and around the lung hilum (Fig. 2E) were milder compared to those near the apex, allowing for easier recognition of the underlying alveolar framework. Within these areas, debris and scattered ruptured macrophages laden with particles were observed in the alveolar spaces (Fig. 2C-left region, enlarged).

The small pure white spots observed macroscopically corresponded to clumps of black pigment in H&E sections (Figs. 2A, 2F). These clumps were observed around the vasculature, subpleural, and within nodular lesions, although they were not prominent within the broncho-vascular bundles of the terminal bronchioles (Fig. 2G). The black pigment in H&E-stained sections represent the TiOL NPs, with the small pure white spots histopathologically identified as clusters of particle-laden macrophages.

In the nodular lesions, we further examined TiOL NPs under polarized light and assessed collagen and elastic fibers, as well as epithelial cell types (Fig. 3). Consistent with previous findings[34,35], TiOL NPs, which were black in H&E staining (Fig. 3A), appeared birefringent under polarized light (Fig. 3B). Masson trichrome staining revealed a marked increase in collagen fibers within the nodules, especially in areas with disrupted alveolar structure, compared to surrounding normal lung tissue (Figs. 3C, S2A-D). In contrast, no increase in elastic fibers[41] was observed in these regions (Fig. 3D).

**Figure 3:**
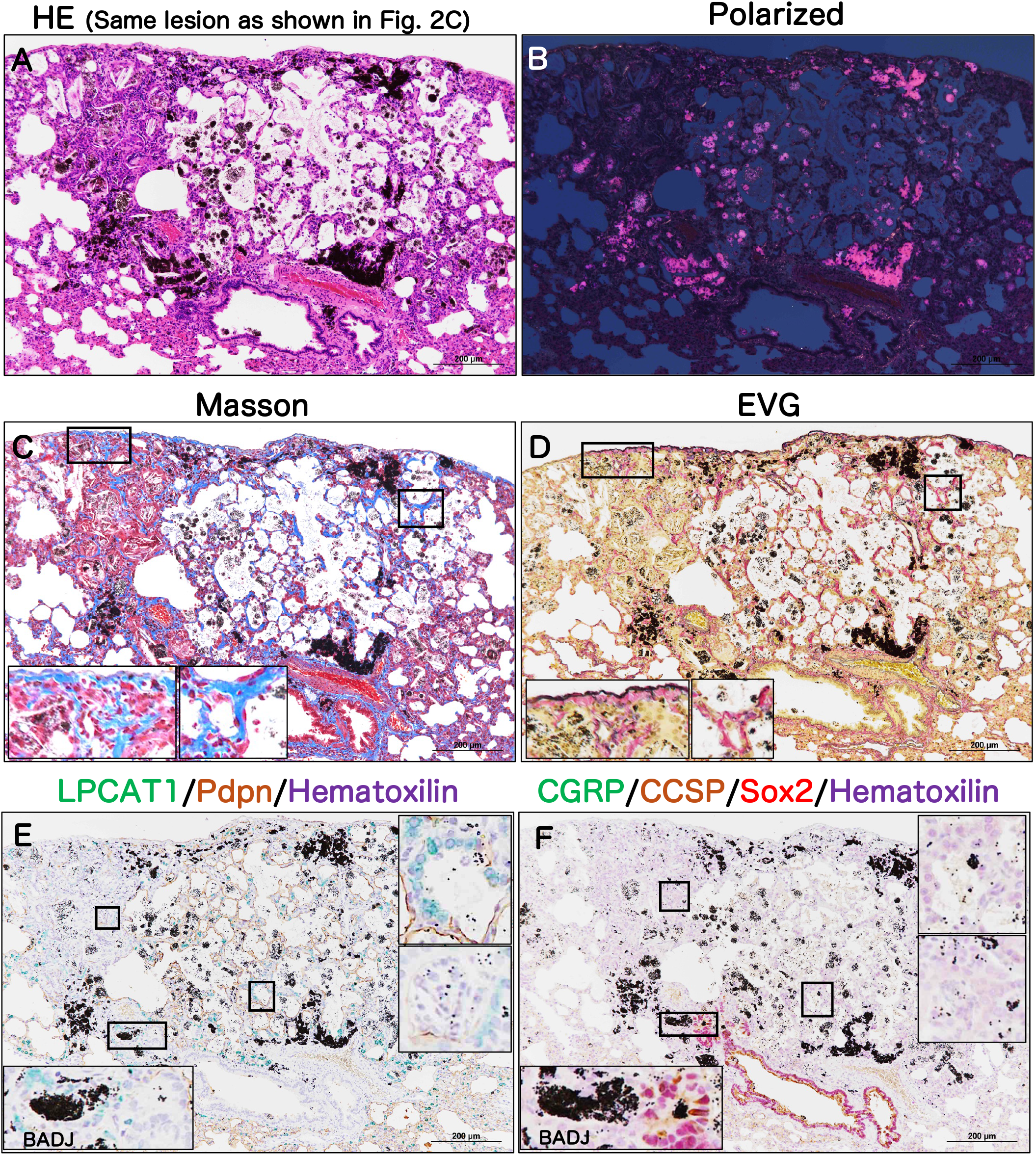
Characteristics of fibrotic pulmonary dust foci (fPDF) in rat lungs after inhalation exposure to TiO_2_ NP (8 mg/m^3^). A: H&E image of the lesion shown in Fig. 2C. The particles show birefringence in polarized light (B). Masson Trichrome staining (C) and EVG staining (D) of the same area show increased collagen fiber deposition within the lesion, but no changes in elastic fibers were observed. Double staining (E) for the AT2 marker LPCAT1 (green in the cytoplasm) and the AT1 marker Pdpn (brown in the cell membrane) and triple staining (F) for the airway neuroendocrine cell marker CGRP (green in the cytoplasm), the airway club cell marker CCSP (brown in the cytoplasm), and the bronchial epithelial lineage marker Sox2 (red in the nucleus). Staining showed that the epithelium comprising the inside of the lesion was of alveolar cell lineage. bar: 200 μm.

To determine the epithelial cell types within the nodules, we performed immunohistochemical (IHC) staining. We used markers for alveolar epithelial lineages (Fig. 3E), LPCAT1 (AT2) and PDPN (AT1), as well as markers for airway epithelial lineages (Fig. 3F), specifically CGRP and Sox2 (neuroendocrine cells) and CCSP and Sox2 (Club cells). Our findings demonstrated that epithelial lineages in the nodular lesions predominantly consisted of alveolar cells.

We further investigated alveolar epithelial lineage markers within the lesions, focusing on transcription factors and markers crucial to alveolar differentiation. Specifically, we examined TTF1, Hop, Sox9, and Sox2, as well as mature alveolar markers LPCAT1 (AT2) and PDPN (AT1) using immunohistochemistry (IHC) (Fig. 4). The analysis showed the presence of TTF1-negative, LPCAT1-positive cells scattered within the lesions (Figs. 4A-C). Additionally, Hop-LPCAT1-double positive cells were identified in areas of alveolar destruction but were absent in regions with milder pathology (Figs. 4D-F). These observations suggest an abnormal differentiation state of alveolar epithelium in fibrotic areas.

**Figure 4:**
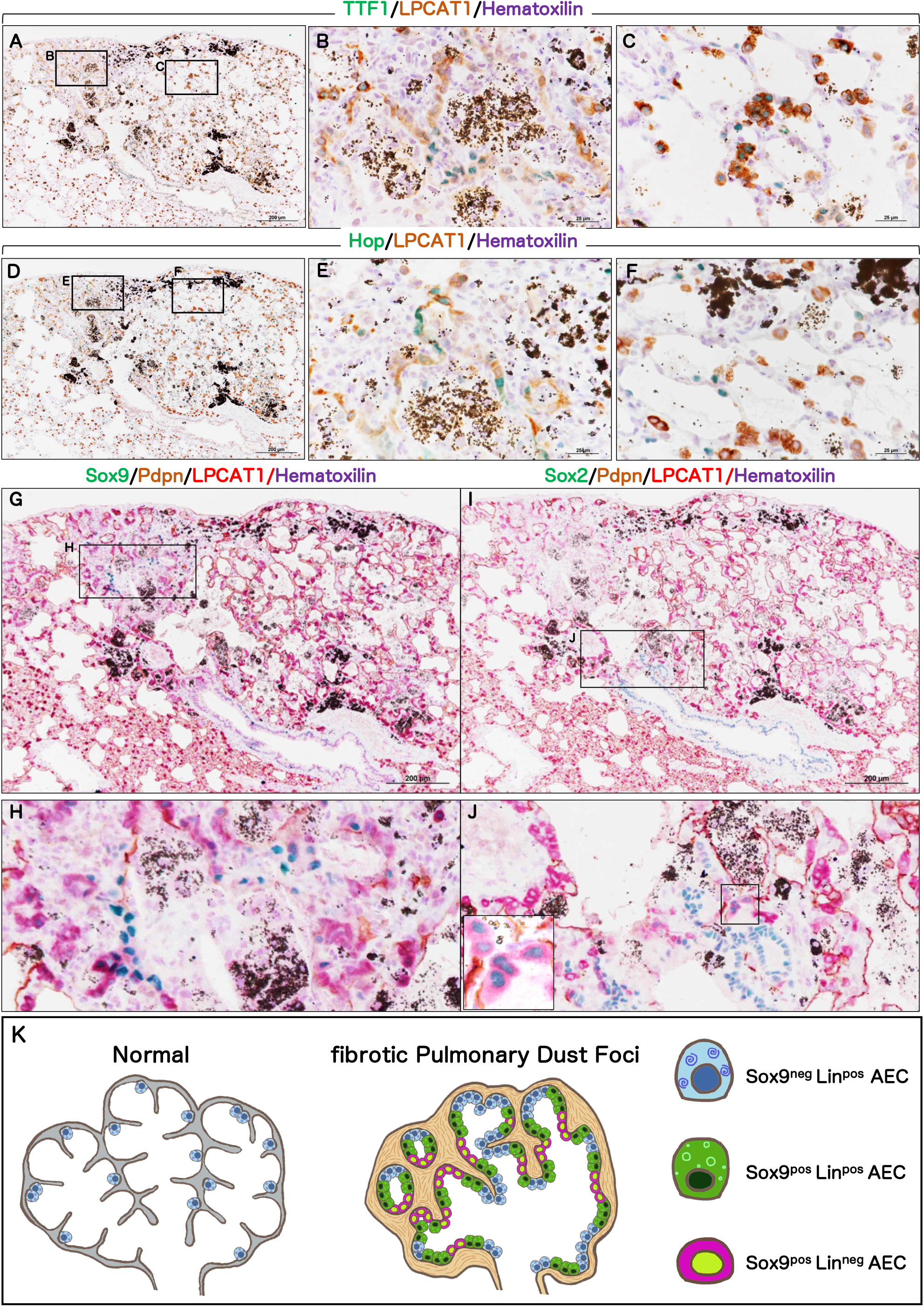
Characteristics of the alveolar epithelial cells in the Fibrotic pulmonary dust foci (fPDF) in rat lungs after inhalation exposure to TiO_2_ NP (8 mg/m^3^). A: Double staining for the AT2 TF marker TTF1 (green in the nucleus) and the AT2 differentiation marker LPCAT1 (brown in the cytoplasm) in the fPDFs. Staining of severe (B) and mild (C) lesions indicate the existence of TTF1-negative AT2 cells. Bar: 200 μm in A, 25 μm in B and C. D: Double staining for the AT1 TF marker Hop (green in the nucleus) and the AT2 differentiation marker LPCAT1 (brown in the cytoplasm) in the fPDFs. Staining of severe (E) and mild (F) lesions indicate the existence of Hop-positive AT2 cells. Bar: 200 μm in D, 25 μm in E and F. G: Triple staining for Sox9 (green in the nucleus), the AT1 differentiation marker Pdpn (brown in the cell membrane), and the AT2 differentiation marker LPCAT1 (red in the cytoplasm) in the fPDFs. H: High magnification of Fig. 4G indicates the presence of Sox9-positive cells in the ducts within the fPDF. I: Triple staining for bronchial epithelial lineage marker Sox2 (green in the nucleus), the AT1 differentiation marker Pdpn (brown in the cell membrane), and the AT2 differentiation marker LPCAT1 (red in the cytoplasm) in the fPDFs. J: High magnification of Fig. 4I indicates the presence of Sox2-positive AT2 cells in the bronchiolar-alveolar duct junction. Bar: 200 μm in G, I. K: Summary of immunostaining results. Lin: Abbreviation for lineage, referring to AT1 or AT2 cells. pos: positive, neg: negative, AEC: alveolar epithelial cell.

Examining Sox9 (Figs. 4G-H) and Sox2 (Figs. 4I-J) revealed that Sox9 was positive in cells within fibrotic regions that were negative for the lineage markers LPCAT1 and PDPN (Fig. 4H). Sox2 was not detected in these fibrotic areas but was present in Sox2-LPCAT1 double-positive cells at the bronchoalveolar junction (Fig. 4J). To summarize (Fig. 4K), normal alveolar epithelium typically consists of Sox9-negative LPCAT1-positive AT2 and PDPN-positive AT1 cells (Sox9^neg^Lin^pos^AEC) in the normal lung alveolar region. In contrast, lesions contained Sox9-positive AT2 or AT1 cells (Sox9^pos^Lin^pos^AEC) as well as Sox9-positive cells lacking both AT2 and AT1 markers (Sox9^pos^Lin^neg^AEC), highlighting an altered differentiation state of alveolar epithelial lineage cells within the fibrotic lesions.

We examined particle deposition and the development of three types of pneumoconiosis lesions (Table 1). Macroscopically, white-gray spots were seen on the lung surface or within the lung tissue. Histologically, these lesions showed marked collagen fiber bundle deposition and inflammation, with infiltration by lymphocytes, neutrophils, mast cells, and plasma cells, alongside cholesterol granulomas containing particle-laden macrophages and multinucleated giant cells. These nodular lesions also exhibited hyperplasia of alveolar epithelium with unstable differentiation. We defined this pattern as "fibrotic pulmonary dust foci (fPDF)." Regarding fPDF progression/severity, mild lesions retained the alveolar framework with fibrous thickening, whereas severe lesions showed alveolar destruction and remodeling, and represented the most advanced pneumoconiosis lesions observed in this study. fPDF occurred in both male and female rats in the 0.5 mg/m³ exposure group, and there was a significant increase in both incidence and severity correlating with exposure concentration, with a higher prevalence in females.

**Table 1:**
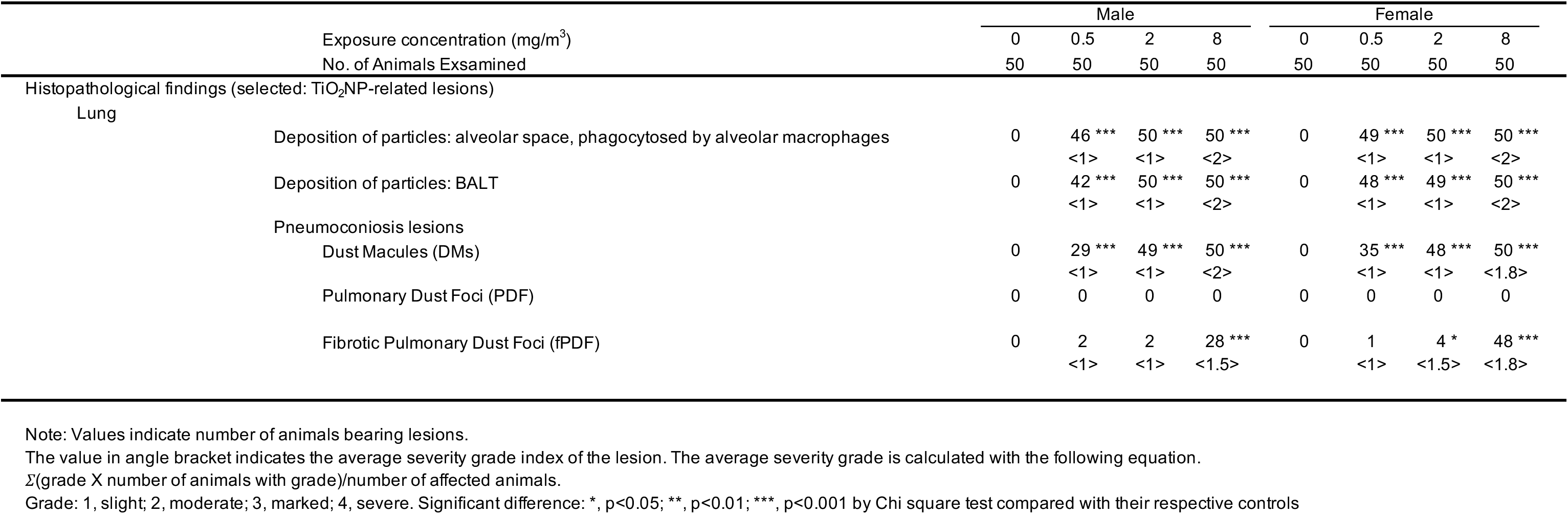
Histopathological findings in the lungs of F344 rats in the carcinogenicity study.

Another lesion type was characterized macroscopically by small, pure white spots and histologically by dense macrophage deposits in the interstitial tissue. Given its similarity to clinical dust-related lesions, we concluded that these lesions were Dust Macules (DMs). DMs were seen in both sexes in the 0.5 mg/m³ exposure group, and also showed a significant exposure-dependent increase in incidence and severity.

We categorized Pulmonary dust foci (PDF) seen in our prior 13-week study[35], fPDFs, and DMs as pneumoconiosis lesions, distinct in nature from simple particle deposits seen in the alveolar spaces or BALT regions. Notably, PDFs were absent in the present study.

### Vascular Characteristics within Fibrotic pulmonary dust foci

We assessed the blood and lymphatic vessel distribution within the fPDFs (Figs. 5, S3). Dual staining with the vascular endothelial marker CD34 and smooth muscle marker αSMA allowed us to trace the arterioles, venules, and capillaries. Compared to surrounding normal tissue, the fPDFs showed a marked reduction in vascular density (Figs. 5A, B). In regions of alveolar destruction/remodeling with severe fibrosis (Fig. 5C) the capillary loss was especially severe, contrasting with areas where alveolar structures were intact (Fig. 5D). Additionally, arterioles and venules were rarely present within the fPDFs. Consistent with these findings, a noticeable reduction in blood-filled vessels was observed in the fPDFs compared to the surrounding normal area. Figure 5E-G shows the blood vessels in an fPDF and the surrounding normal tissue of a rat that died at week 101. These findings indicate that the capillary network is poorly developed in the severely fibrotic areas where alveolar destruction/remodeling is progressing in fPDFs.

**Figure 5:**
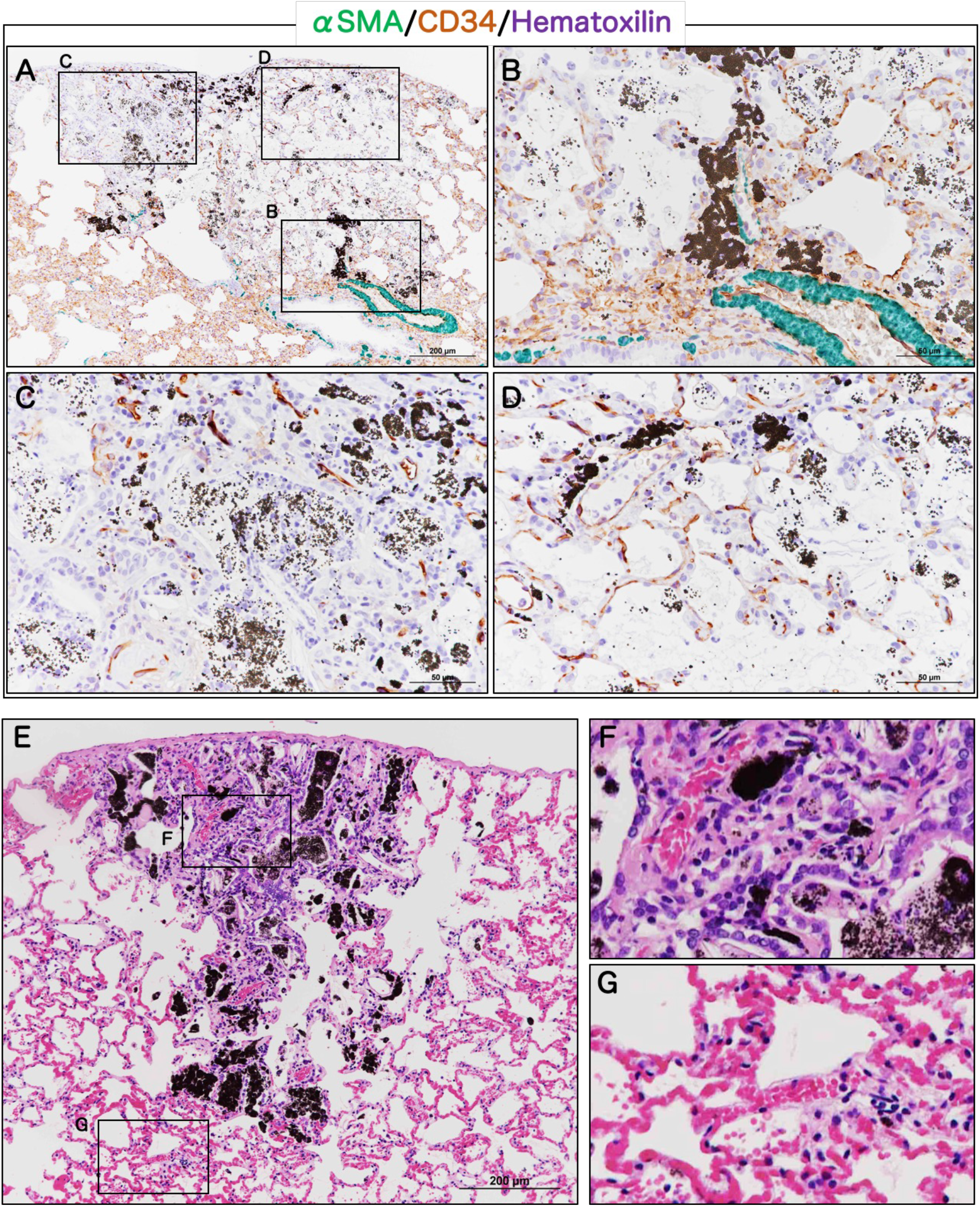
Blood Vascular Characteristics of Fibrotic pulmonary dust foci (fPDF) in rat lungs after inhalation exposure to TiO_2_ NP (8 mg/m^3^). A: Double staining for the smooth muscle cell marker αSMA (green in the cytoplasm) and the vascular endothelial cell marker CD34 (brown in the cell membrane) in the fPDFs. B-D: High magnification of A indicates that the capillary network is poorly developed, especially in the severely fibrotic areas where alveolar destruction/remodeling is progressing in fPDF. Bar: 200 μm in A, 50 μm in B-D. E: H&E staining image of the lung tissue of a rat in the 8mg/m3 exposure group that died at week 101. F-G: High magnification of E indicates a marked accumulation of red blood cells (RBCs) in the capillaries of normal surrounding tissue and a loss of RBCs in the fPDF. Bar: 200 μm in E.

Lymphatic vessel assessment, using Prox1 (nuclear marker) and VEGFR3 (membrane marker), revealed that lymphatic elongations reached the areas surrounding the fPDF-normal tissue boundaries (Figs. S3A-C). Lymphatic vessels, absent in normal alveolar areas, extended into the interstitium within the lesions (Figs. S3D-E). These results highlight an imbalance in vascular distribution within the fPDFs, a defining feature of the pathological components forming these regions.

### Characteristics of Interstitial Macrophages

To explore the distribution of macrophages that had phagocytosed TiOL NPs, we used the nuclear marker PU.1 and the cytoplasmic marker CD68 to identify macrophages, and Pdpn as an AT1 marker to define alveolar frameworks and distinguish alveolar airspaces from interstitial regions. In fPDFs, macrophages were confined to granulomas containing particle-laden macrophages and multinucleated giant cells. In contrast to fPDFs, free macrophages containing TiOL NPs were frequently located in DMs in the subpleural and perivascular interstitial regions surrounding arterioles and venules (Figs. 6A, B). Fewer macrophages were observed in bronchus-associated lymphoid tissue (BALT) (Fig. 6C) and around interstitial broncho-vascular bundles (Fig. 6D). Notably, macrophages in the alveolar airspaces exhibited a marked tendency toward destruction while interstitial macrophages loaded with TiOL NPs showed no signs of destruction. This suggests that the particle toxicity varied in these different macrophage populations, possibly due to attempted destruction of the phagocytosed TiOL NPs in order to contain "the infection" by the alveolar macrophages and simple removal of the TiOL NPs by the interstitial macrophages.

**Figure 6:**
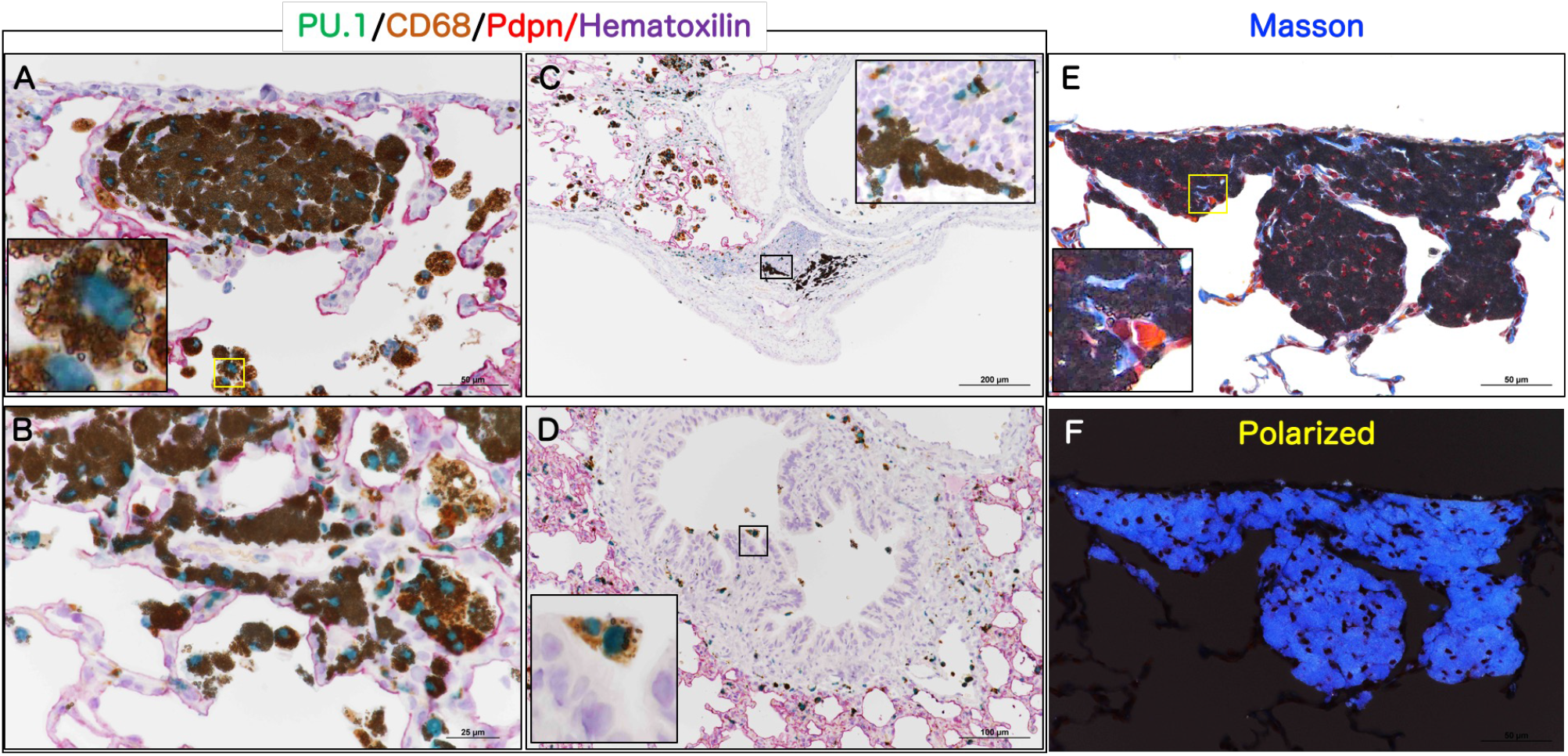
Characteristics of Interstitial Macrophages A-D: Triple staining for the myeloid lineage marker PU.1 (green in the nucleus), the macrophage marker CD68 (brown in the cytoplasm), and the AT1 differentiation marker Pdpn (red in the cell membrane) in the subpleural interstitium (A, bar: 50 μm), perivascular interstitium (B, bar: 25 μm), BALT (C, bar: 200 μm), and broncho-vascular bundle (D, bar: 100 μm) of a rat lung exposed to 8 mg/m^3^ TiO_2_ NPs. E: Masson Trichrome staining of a dust macule. F: Polarized image in E. Bar: 50 μm in E-F.

To assess potential fibrosis associated with DM, we performed Masson Trichrome staining to evaluate collagen fibers around macrophages (Figs. 6E, F). In contrast to the highly fibrotic fPDF lesions, collagen fiber levels in DMs were similar to those in normal interstitial tissue, with no evidence of fibrotic change. These data indicate that the toxicity of TiOL NPs to interstitial macrophages is limited, and there is no excessive deposition of collagen within DMs.

### Continuity Between Pneumoconiosis Lesions and Pulmonary Tumors

Figs. S4A-D show typical macroscopic and microscopic images of bronchiolo-alveolar carcinoma in the TiOL NPs -exposure group. This carcinoma was identified in areas with few white-gray spots (Fig. S4A, B), and no histopathological continuity was observed between the pneumoconiosis lesions, both fPDFs and DMs, and the carcinoma (Figs. S4C, E). Moreover, Particle-laden macrophages were also rare within the central position of the tumor (Fig. S4D) and along its margins (Fig. S4F).

Futhermore, lung tumors in the exposed groups displayed histopathological features—such as tumor cell growth patterns and stromal architecture—similar to those observed in spontaneous, age-related lung tumors. These findings suggest that pneumoconiosis and the proliferative changes of the alveolar epithelium, which are components of pneumoconiosis, were not directly related to the development of lung tumours in this study.

## Discussion

We previously investigated the carcinogenicity of titanium dioxide nanoparticles (TiO_2_ NPs) by exposing F344/DuCrlCrlj rats to TiO_2_ NP aerosols at concentrations of 0, 0.5, 2, and 8 mg/m^3^ for 6 h/day, 5 days/week for 104 weeks using a whole-body inhalation exposure system. We concluded that there was no clear evidence of carcinogenicity in male or female rats[40]. We also conducted a medium-term (26 weeks) whole-body inhalation exposure study with mice using the same TiO_2_ NPs we used in the 2-year rat study. We found no evidence of carcinogenicity in the lungs or other organs of mice exposed to 2, 8, or 32 mg/m^3^ TiO_2_ NPs[34]. In the present study, we re-evaluated lung pathology in rats after two years of inhalation exposure to TiOL NP to assess pneumoconiosis pathology. Rats exposed to TiOL NPs developed white lung lesions, primarily in subpleural peripheral areas. Microscopically, two key pneumoconiosis lesion types were identified (Fig. 7): (1) Fibrotic pulmonary dust foci (fPDF), characterized by severe collagen deposition, wide-spectrum inflammatory cell infiltration, and AEC hyperplasia, and (2) Dust Macule (DM), marked by particle-laden macrophage accumulation in the interstitium not accompanied by collagen fiber deposition, infiltration of other inflammatory cells infiltration, or AEC hyperplasia.

**Figure 7:**
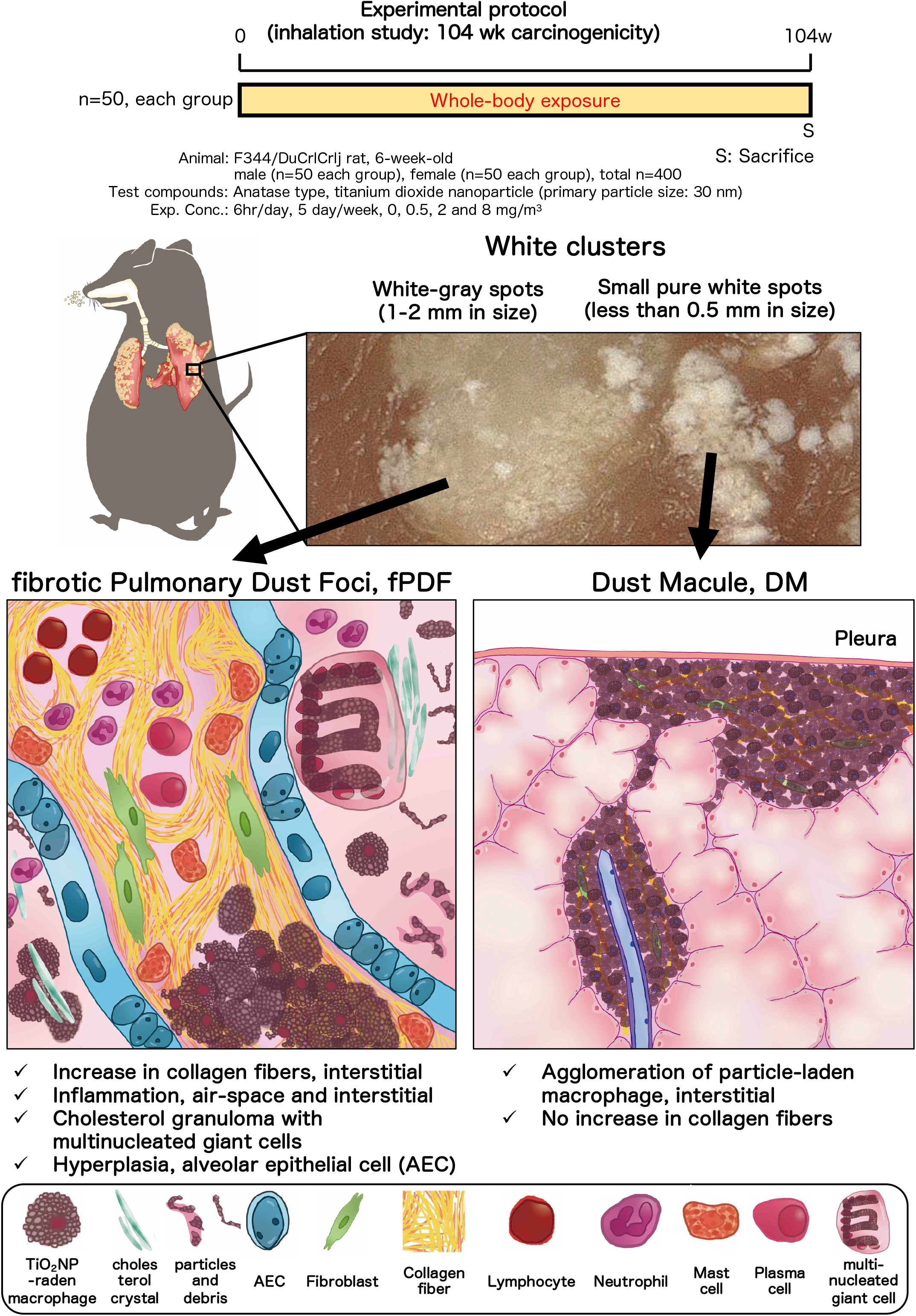
Graphical abstract in this study.

Detailed immunohistochemical analysis highlighted disrupted alveolar epithelial cell differentiation within the fPDF. Notably, fPDF exhibited significantly reduced capillary density, impacting blood and lymphatic vessel distribution. Despite the high dose of TiOL NPs exposure, the tumors observed in exposed rats showed no histological connection to pneumoconiosis-related lesions, appearing similar to spontaneous age-related tumor histology. Based on the previous assessment of carcinogenesis and the assessment of pneumoconiosis in the present study, we conclude that 2 years exposure to TiOL NP causes advanced pneumoconiosis in rat lungs but does not directly promote lung tumor development.

We previously reported a characteristic early pneumoconiosis lesion that we defined as pulmonary dust foci (PDF) following 13-weeks inhalation exposure of rats to TiOL NPs[35]. Histopathologically, PDFs were characterized by aggregates of particle-laden macrophages within the alveolar airspace, surrounded by hyperplasia of type II alveolar epithelial cells (AT2), without collagen fiber deposition in the alveolar septa. Based on these findings, we concluded that PDFs represent an early pneumoconiosis lesion in rats. In the present study, we found that fPDFs shared anatomical location with PDFs identified in the 13-week inhalation study but displayed more diverse inflammatory spectra, including collagen fiber deposition. This suggests that fPDFs represent advanced forms of PDFs. Macrophages laden with particles within the airspace were a critical component of pneumoconiosis lesions. However, compared to the 13-week inhalation study, these air-space macrophages appeared more fragile in the 2-year carcinogenicity study, leading to cell death and the presence of fragmented macrophages and debris. These findings suggest that the vulnerability of macrophages in the airspace is a significant and clear difference between the early lesions (PDF) and advanced lesions (fPDF) observed in the present study. The alveolar epithelium, a key component of the PDF-fPDF axis, exhibited persistent abnormalities. In PDFs, AT2 cells showed increased γH2AX expression[35], while fPDFs exhibited defects in the differentiation state of alveolar lineage cells. However, there was no clear evidence linking these findings to the bronchiolo-alveolar hyperplasia-adenoma-carcinoma axis, a pathological hallmark of rat lung carcinogenesis. In addition, in contrast to pneumoconiosis lesions, there was no clear increase in carcinogenicity with increasing TiOL NP exposure levels[40]. These results suggest that lesions in the PDF-fPDF axis and DNA damage in AEC caused by TiOL NP exposure in rats are not reliable and sensitive markers for predicting pulmonary carcinogenicity. They also highlight the need to reevaluate the proposed Adverse Outcome Pathway for TiOL NP inhalation exposure[42].

In workers, a wide variety of pathological responses to inhaled dust have been reported[10,43]. Specifically, individuals exposed to mixed dust containing various ratios of crystalline silica and silicates typically exhibit three types of pathological lesions in the interstitium surrounding the respiratory bronchioles (also referred to as the broncho-vascular bundle, BVB). These include dust macules (DMs), mixed-dust fibrotic lesions (MDFs), and silicotic nodules (SNs).When the crystalline silica/silicate concentration in the mixed dust is extremely low, at 1% or less, and the dust is inert or has weak fibrotic effects, it forms an impalpable interstitial lesion known as a dust macule (DM). These are composed of aggregates of dust-laden macrophages. In contrast, MDFs arise from exposure to mixed dust with a crystalline silica/silicate concentration of 20% or less. These palpable stellate lesions with irregular outlines vary in the degree of collagenization. SNs result from mixed dust with crystalline silica/silicate concentrations exceeding 20%. These lesions consist of highly fibrotic, glassy structures, displaying layered concentric or irregular arrangements, and contain minimal cellular components. In summary, mixed dust inhaled by workers leads to solitary clustered lesions in the interstitium surrounding the respiratory bronchioles.

These lesions represent a spectrum of interstitial abnormalities, ranging from non-fibrotic DMs to mildly fibrotic MDFs and highly fibrotic SNs, with varying severity.

The present study demonstrated that DMs can develop in rodents, but their onset requires an extended period. DMs were not observed in a 13-week inhalation study, emphasizing the time-dependent nature of this lesion. Although DMs in humans and rats share similar cellular compositions, their preferred locations differ. In humans, DMs typically form around BVBs, while in rats, they are more common near subpleural regions. This disparity may reflect anatomical differences, such as the shorter airway length and closer proximity to the pleura in rodents, as well as differences in lymphatic pathways. Notably, MDFs did not develop in rats, even after two years inhalation exposure to TiOL NP. This suggests that interstitially located TiOL NPs or the macrophages that engulf them have low fibrogenic potential. These findings are consistent with the classification of TiOL NP as poorly soluble, low toxicity particle[44]. In this study, the most severe pneumoconiosis-related lesions in rats were fPDFs. These lesions are distinct from the interstitial DM-MDF-SN axis observed in humans. The fPDF lesions began as PDF lesions during the subacute phase and gradually developed into fPDFs, marked by increased fibrosis. The findings suggest that persistent inflammatory responses in the air spaces play a key role in disease progression. The pulmonary changes observed in rats exposed to TiOL NPs are consistent with prior findings from long-term inhalation studies in rats[45]. These results also align with reports indicating that rats have limited capacity for interstitial migration of dust particles, reflecting a species-specific response[46,47]. Previous studies have reported clustering lesions of particle-laden macrophages in mice, rats, and hamsters exposed to nuisance and relatively nonfibrogenic dusts[48–52]. Green *et al* described the development of DM after inhalation of such particles, including coal mine dust[53]. However, these lesions were primarily located in the air spaces and were qualitatively different from the interstitial DM found in humans. Thus far, there have been no reports of DMs in rats resembling the interstitial DM-MDF-SN axis observed in humans. Our detailed histopathological re-analysis has identified the development of DMs in rats exposed to TiOL NPs that are similar to DMs found in workers.

A key limitation of this study is the two-year observation period, during which all animals were sacrificed and evaluated at death or the two-year endpoint. In contrast, previous studies have extended observations to 2.5 years or longer lifetime durations. Thus, our results cannot exclude the possibility of carcinogenesis or progression to MDF occurring with longer exposure periods.

## Conclusions

We re-evaluated the pulmonary lesions caused by two years of inhalation exposure to anatase-type titanium dioxide nanoparticles (TiO_2_ NP) in F344 rats from the perspective of pneumoconiosis pathology. The findings revealed two types of pneumoconiosis lesions: fibrotic pulmonary dust foci (fPDF) and dust macules (DM).

The fPDF lesions were progressive forms of PDF, previously observed and defined in our 13-week inhalation exposure study. fPDF lesions primarily featured increased collagen fiber bundles and a broader spectrum of inflammatory responses. They were also associated with abnormal differentiation of alveolar epithelial cell lineages and vascular structures. In contrast, DM lesions, which are frequently observed in workers, were interstitial lesions characterized by clusters of particle-laden macrophages. These lesions displayed no increase in collagen fibers and lacked diverse inflammatory cell infiltration. Neither fPDF or DM lesion types exhibited direct continuity with the bronchiolo-alveolar hyperplasia-adenoma-carcinoma sequence, strongly suggesting that in the current study pneumoconiosis did not lead to tumorigenesis. In addition, the tumors that developed in the TiO_2_ NP exposed rats resembled spontaneous, age-related neoplasms, further supporting the premise that in the present study development of pneumoconiosis lesions did not lead to tumorigenesis.

In conclusion, long-term inhalation of titanium dioxide nanoparticles (TiO_2_ NP) in rats primarily induced pulmonary lesions originating from the PDF-fPDF axis. These lesions began in the air spaces and spread into the interstitium, with most effects localized to the alveolar regions. Additionally, DM lesions similar to those found in workers were also identified.

## Material and method Materials

Lung tissue samples obtained from a two-year inhalation carcinogenicity study were used as materials. Detailed information on the carcinogenicity study has been described in a previous report [40]. Briefly, Anatase-type TiO_2_ NP was purchased from Tayca co. (purity of > 97.9%; Lot No. 6545, primary particle size: 30 nm). F344/DuCrlCrlj (SPF) rats (Charles River Japan Inc.; Kanagawa, Japan) of each sex were exposed to TiO_2_ NP aerosol for 104 weeks (6 hr/day, 5 days/week exposure cycles) at target concentrations of 0, 0.5, 2, and 8 mg/m^3^. Animals were sacrificed after the end of exposure on 5 separate days. The left lung was fixed by 10% buffered formalin injection, followed by immersion fixation. The right lung was directly immersion-fixed. After fixation for approximately 2-7 days, each lobe of the lung was sectioned for microscopic observation (Fig. S1) and prepared for paraffin embedding.

### Histopathological analysis

Serial tissue sections were cut from formalin-fixed paraffin-embedded lung specimens, and the first sections (2-μm thick) was stained with H&E for histological examination and the remaining sections were used for immunohistochemical analysis. The histopathological findings in this study were determined by two certified pathologists from the Japanese Society of Toxicologic Pathology. Diagnostic terminology was based on the International Harmonization of Nomenclature and Diagnostic Criteria for Lesions in Rats and Mice (INHAND) [54] and the textbook for human pneumoconiosis diagnosis[4,5]. Non-neoplastic lesion was evaluated for its severity and scored on a scale of “slight” to “severe” with reference to the criteria by Shackelford et al[55]. The section was observed under an optical microscope ECLIPSE Ni (Nikon Corp., Tokyo, Japan) or BZ-X810 (Keyence, Osaka, Japan) or scanned with the VS120 virtual microscopy slide scanning system (Olympus, Tokyo, Japan).

### Masson’s trichrome staining

The slides were deparaffinized, washed with water, and then reacted with a mixture of 10% trichloroacetic acid and 10% potassium dichromate for 60 min at room temperature. The specimens were then washed with water and stained with Weigert’s iron hematoxylin solution (C.I.75290, Merck-Millipore) for 10 min at room temperature (RT). The slides were then successively stained with 0.8% orange G solution (C.I.16230, Merck-Millipore) for 10 min at RT, Ponceau (C.I.14700, FUJIFILM-Wako Pure Chemical Corp., Osaka, Japan) acid fuchsin (C.I.42685, Merck-Millipore) azophloxine (C.I.18050, Chroma Germany GmbH, Augsburg, Germany) mixture for 40 min at RT, 2.5% phosphotungstic acid for 10 min at RT, and blue aniline solution (C.I.42755, Chroma Germany GmbH) for 10 min at RT. Between each staining the slides were washed lightly with 1% acetic acid water and then dehydration, permeabilization, and sealing were performed.

### Elastica Van Gieson staining

Details of the multiple staining method have been described previously[35]. Briefly, the slides were deparaffinized, washed with water, reacted with Maeda Modified Resorcinol-Fuchsin Staining Solution (Mutoh Chemical, Part No. 40321, Japan) for 30 min at RT, and rinsed with 100% ethanol to remove excess stain. The slides were washed with running water, stained with Weigelt’s iron hematoxylin solution (C.I.75290, Merck-Millipore, US) for 10 min at RT, and washed with running water for 10 min. The slides were then reacted with 1% Sirius red solution (Mutoh Chemical, Part No. 33061, Japan) for 3–5 min at RT, washed with water, dehydrated with 90%-100% ethanol, permeabilized, and sealed.

### Immunohistological multiple staining analyses

Details of the Immunohistological multiple staining method have been described previously[37,56]. Briefly, lung tissue sections were deparaffinized with xylene, then hydrated through a graded ethanol series, and incubated with 0.3% hydrogen peroxide for 10 min to block endogenous peroxidase activity. Slides were then incubated with 10% normal serum at RT for 10 min to block background staining, and then incubated for 2 h at RT with the first primary antibody (see Table S1). After washing with PBS, the slides were incubated with histofine simple stain ratMAX-PO(MULTI) (414191, Nichirei, Tokyo, Japan) for 30 min at RT. After washing with PBS, slides were incubated with DAB EqV Peroxidase Substrate Kit, ImmPACT (SK-4103, Vector laboratories) for 2-5 min at RT. Importantly, after washing with dH_2_O after color detection, the sections were treated with citrate buffer at 98 °C for 30 min before incubation with the next primary antibody to denature the antibodies already bound to the section. This procedure was repeated for the second and then the third primary antibody. HighDef red IHC chromogen (ADI-950-142, Enzo Life Sciences, Inc., Farmingdale, NY, USA) was used for the second coloration and Histogreen chromogen (AYS-E109, Cosmo Bio, Tokyo, Japan) for the third coloration. Coloration was followed by hematoxylin staining for 30-45 s. The slides were then processed for light microscopy.

### Statistical analysis

Except for the incidence and integrity of histopathological lesions, the data comparisons among multiple groups were performed as follows: when normal distribution and homogeneous variance were observed in samples without sex differences, a one–way ANOVA was used to compare the exposure and control groups. When the one-way ANOVA was significant, Dunnett’s multiple comparisons test was used to compare the control and exposure groups. If variances were significantly different, the control and exposure groups were evaluated using Kruskal-Wallis non-parametric analysis of variance. If the Kruskal-Wallis analysis was significant, the control and exposure groups were compared using Dunn’s test. Those statistical analyses were performed with GraphPad Prism 5 (GraphPad Software). The incidences and integrity of lesions were analyzed by the chi-square test using GraphPad Prism 5 (GraphPad Software). Statistical significance was set at pL<L0.05.

## Abbreviations

αSMA: α-smooth muscle actin
AEC: alveolar epithelial cell
AT1: alveolar epithelial type 1 cell
AT2: alveolar epithelial type 2 cell
BALT: bronchus-associated lymphoid tissue
BVB: broncho-vascular bundle
CCSP: club cell secretory protein
CGRP: calcitonin gene-related peptide
DM: dust macule
fPDF: Fibrotic pulmonary dust foci
γH2AX: Phosphorylation of the Ser-139 residue of the histone variant
H2AX HE: hematoxylin and eosin
Hop: homeodomain-only protein
IHC: immunohistochemistry
INHAND: International Harmonization of Nomenclature and Diagnostic Criteria for Lesions in Rats and Mice
Lin: Lineage
LPCAT1: lysophosphatidylcholine acyltransferase 1
MDF: Mixed-dust fibrotic lesion
OECD: Organization for the Economic Co-operation and Development
PDF: pulmonary dust foci
Pdpn: podoplanin
Prox1: Prospero homeobox protein-1
PU.1: Spi-1 Proto-Oncogene
SEM: scanning electron microscope SN: Silicotic nodule
Sox2: SRY-Box Transcription Factor 2
Sox9: SRY-Box Transcription Factor
9 SPF: Specific Pathogen Free
SUR: tissue surrounding a lesion
TiO_2_ NPs: titanium dioxide nanoparticles
TTF1: Thyroid Transcription Factor 1
VEGFR3: Vascular endothelial growth factor receptor 3

## Declarations

### Ethics approval and consent to participate

All animals were treated humanely, and all procedures were performed in compliance with the Animal Experiment Committee of the Japan Bioassay Research Center.

## Consent for publication

All authors gave their consent for publication of this manuscript.

## Availability of data and materials

The datasets used and analyzed during the current study are available from the corresponding authors on reasonable request.

## Competing interests

The authors declare that they have no competing interests.

## Acknowledgments

We wish to thank Dr. David B. Alexander of Nanotoxicology project, Nagoya City University Graduate School of Medicine for his insightful comments and English editing. We would also like to thank Misae Saito for her assistance with the pathological experiments, Aya Umeda for drawing the fig.4K, and Akane Yoshida for drawing the fig.7. Figs. 1G, S4B-F, and S3D in this paper have already been published in the following references.

## Reference 1 (Figs. 1G, S4B-F)

Kasai T, Hirai S, Furukawa Y, Misumi K, Takeda T, Goto Y, Takanobu K, Yoneyama K, Yamano S, Senoh H, Umeda Y. Lung carcinogenicity by whole body inhalation exposure to Anatase-type Nano- titanium dioxide in rats. J Toxicol Sci. 2024;49(8):359-383. doi: 10.2131/jts.49.359. PMID: 39098045.

## Reference 2 (Fig. S3D)

Takeda T, Yamano S, Goto Y, Hirai S, Furukawa Y, Kikuchi Y, Misumi K, Suzuki M, Takanobu K, Senoh H, Saito M, Kondo H, Daghlian G, Hong YK, Yoshimatsu Y, Hirashima M, Kobashi Y, Okamoto K, Kishimoto T, Umeda Y. Dose-response relationship of pulmonary disorders by inhalation exposure to cross-linked water-soluble acrylic acid polymers in F344 rats. Part Fibre Toxicol. 2022 Apr 8;19(1):27. doi: 10.1186/s12989-022-00468-9. Erratum in: Part Fibre Toxicol. 2022 May 13;19(1):35. doi: 10.1186/s12989-022-00475-w. PMID: 35395797; PMCID: PMC8994297.

## Funding

This research was financially supported by a grant-in-aid from the Japan Organization of Occupational Health and Safety.

## Author information

National Institute of Occupational Safety and Health, Japan Organization of Occupational Health and Safety, Fujisawa, Kanagawa 251-0015, Japan Shotaro Yamano, Yumi Umeda

## Contributions

S.Y. and Y.U. performed the experiments, analyzed the data, histopathological diagnoses, drafted and revised the manuscript. All authors approved the manuscript as submitted.

## Corresponding authors

Correspondence to Shotaro Yamano or Yumi Umeda

## Supplementary Information

**Additional file 1: Figure S1.** Method of cutting out the lung organ. The left lung of the rat lung is inflated with formalin and then cut into two sections. The right lung is not inflated with formalin and the accessory lobe is woven into the back side and cut into one section, and a total of three cross-sections are made into formalin-fixed paraffin embed blocks.

**Additional file 2: Figure S2:** Masson Trichrome staining of the Fibrotic pulmonary dust foci (fPDF) in rat a lung after inhalation exposure to TiO_2_ NP (8 mg/m^3^). Serial tissue staining of a fPDF with marked destruction/remodeling of alveolar structure accompanied by an increase in collagen bundles. H&E (A, B) and Masson Trichrome (C, D). Bar: 100 μm in A-B, and 25 μm in C-D.

**Additional file 3: Figure S3:** Lymphatic Vascular Characteristics of Fibrotic pulmonary dust foci (fPDF) in a rat lung after inhalation exposure to TiO_2_ NP (8 mg/m^3^). A: Double staining for the lymphatic endothelial cell lineage marker Prox1 (green in the nucleus) and the lymphatic endothelial cell marker VEGFR3 (brown in the cell membrane) in a fPDF. B-C: High magnification of A indicates that lymphatic vessel elongations reached the areas surrounding the fPDF-normal tissue boundaries. Bar: 200 μm in A, 25 μm in B-C. D-E: Summary of immunostaining results. TB: Terminal bronchiole, AD: Alveolar duct, AS: Alveolar sac, A: Alveori, Ar: Arteriole, Vn: Venule, Ly: Lymphatic vessels.

**Additional file 4: Figure S4:** Representative macroscopic and microscopic photographs of a bronchiolo-alveolar carcinoma in a male rat lung after inhalation exposure to TiO_2_ NP (8 mg/m^3^). A: Representative macroscopic photographs of a bronchiolo-alveolar carcinoma in a male rat left lung after inhalation exposure to TiO_2_ NPs (8 mg/m^3^). Bar: 1 cm. B: Loupe image of H&E-stained lung sections of A. Bar: 5 mm. C-F: High magnification of central (C-D) and peripheral (E-F) regions of the bronchiolo-alveolar carcinoma. Bar: 250 μm in C, E, 50 μm in D-F.

**Additional file 5: Table S1:** List of primary antibodies used in this study.

## Notes

### Competing Interest Statement

The authors have declared no competing interest.

