## Supplementary figures and images for "Fibrotic pulmonary dust foci is an advanced pneumoconiosis lesion in rats induced by titanium dioxide nanoparticles in a 2-year inhalation study"

### supplemental figures

Figure S1

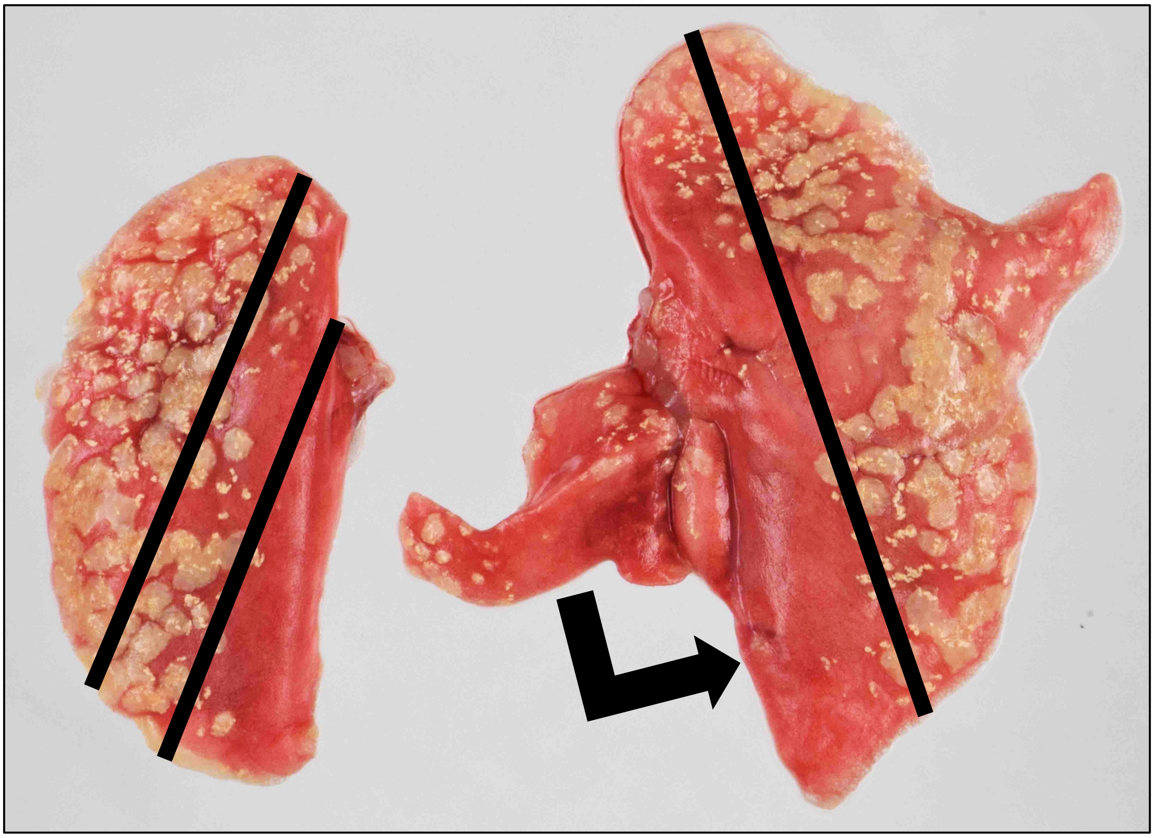

Figure S2

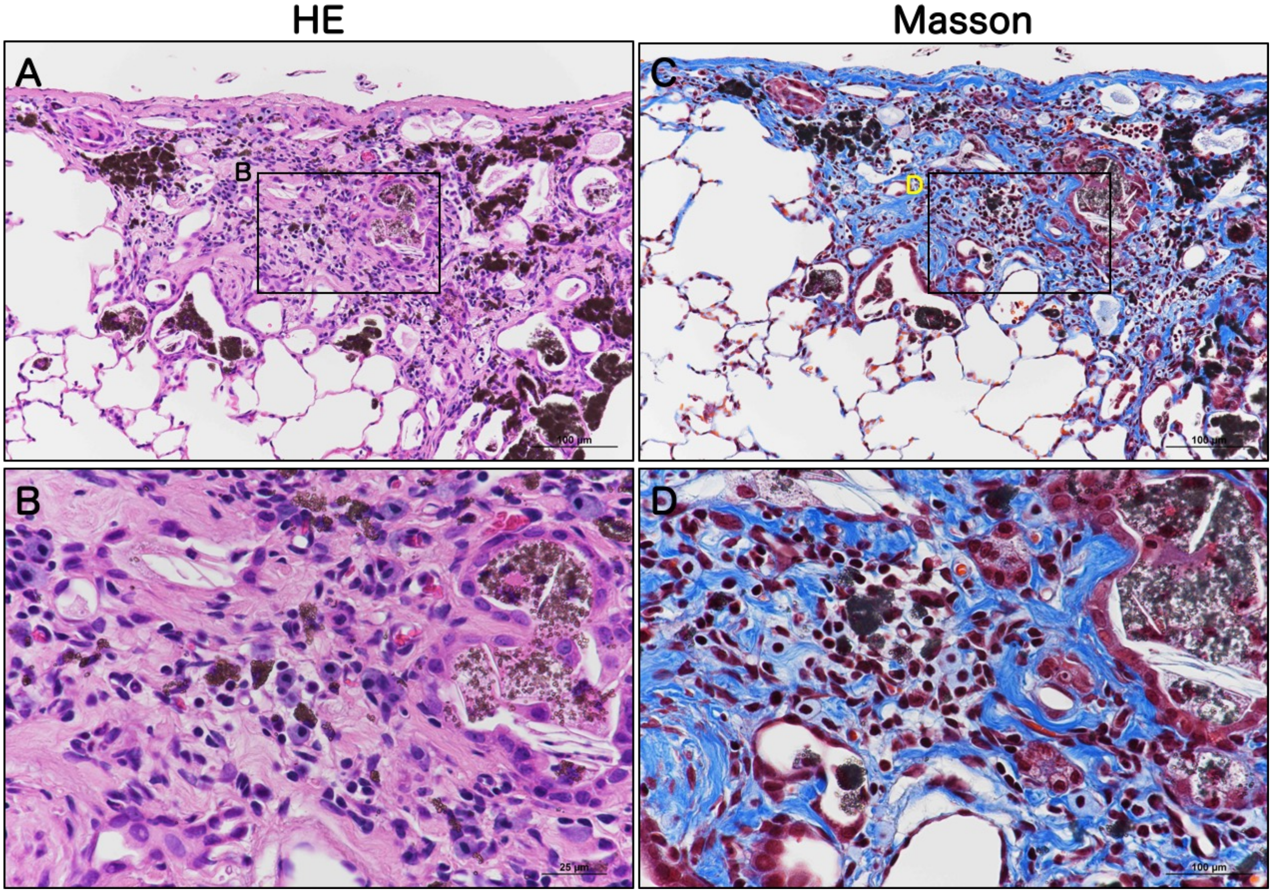

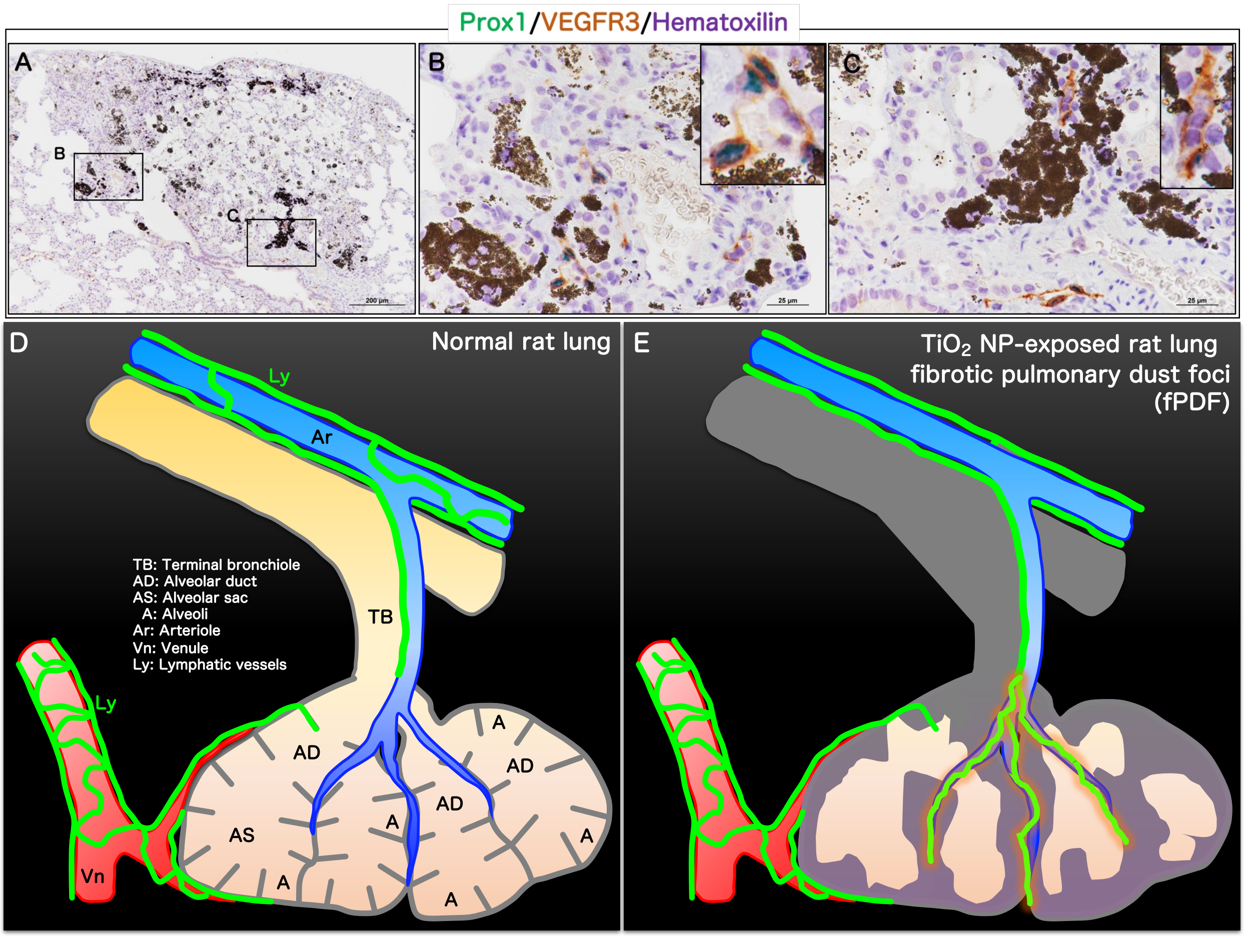

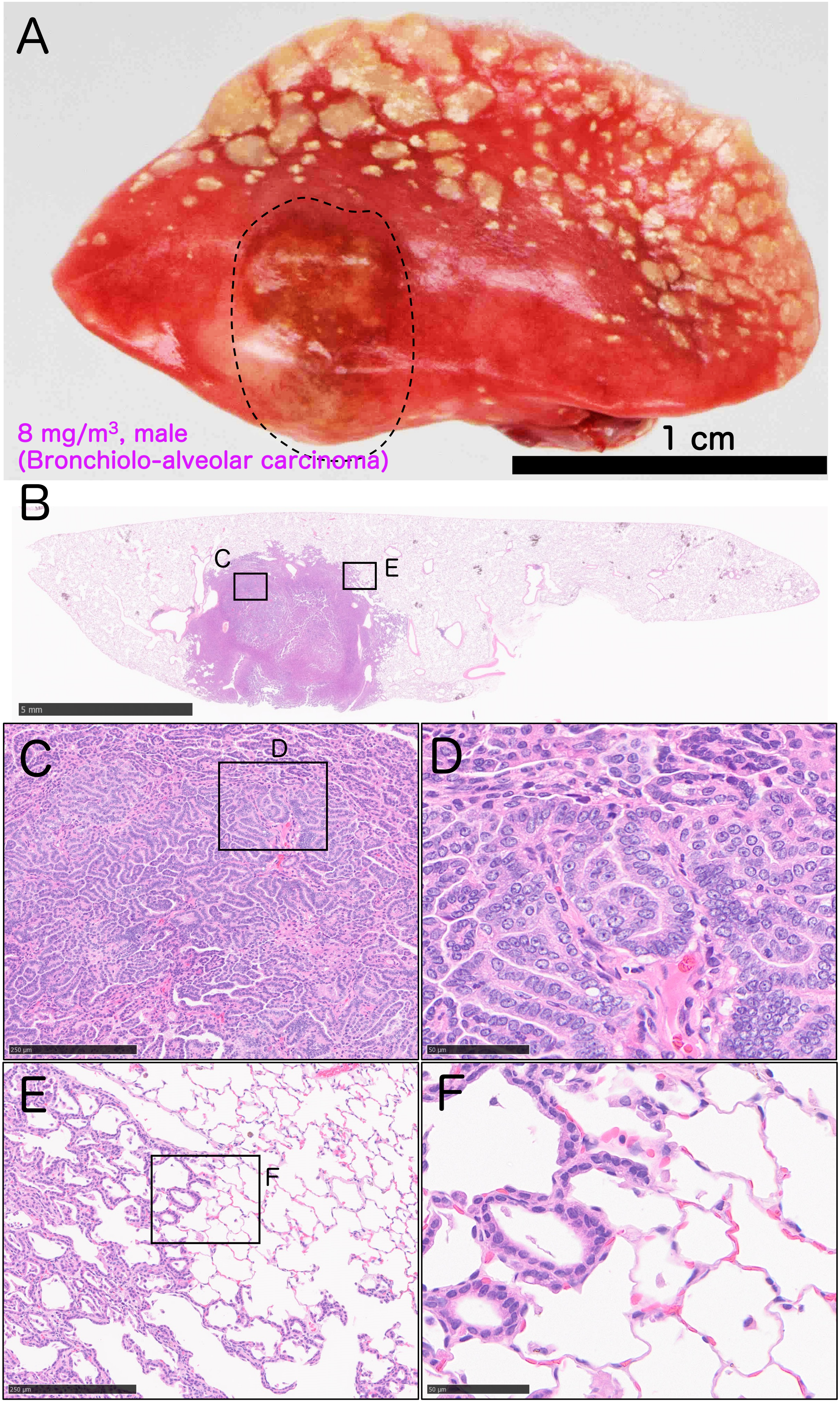
