## Supplementary material for "Fibrotic pulmonary dust foci is an advanced pneumoconiosis lesion in rats induced by titanium dioxide nanoparticles in a 2-year inhalation study": Table s1

Table S1　A list of all primary antibodies.

| 1st Antibodies | product company | ID | host | clonarity | dilution |
|---|---|---|---|---|---|
| CCSP | Sevenhillis bioreagents | WRAB-3950 | rabbit | poly | 1:20000 |
| CGRP | abcam | ab36001 | goat | poly | 1:1000 |
| Sox2 | abcam | ab93689 | rabbit | mono | 1:100 |
| Sox9 | abcam | ab185966 | rabbit | mono | 1:200 |
| Hop | santacruz | sc-398703 | mouse | mono | 1:1000 |
| TTF1 | DAKO | M3575 | mouse | mono | 1:100 |
| LPCAT1 | Proteintech | 66044-1-Ig | mouse | mono | 1:1000 |
| αSMA | abcam | ab5694 | rabbit | poly | 1:500 |
| PU.1 | santacruz | sc-390659 | mouse | mono | 1:100 |
| Prox1 | R&D | AF2727 | goat | poly | 1:100 |
| VEGFR3 | R&D | AF743 | goat | poly | 1:250 |
| CD34 | R&D | AF4117 | goat | poly | 1:1000 |
| RT1-40 (podoplanin) | terracebiotech | TB-11ART1-40 | mouse | mono | 1:100 |
| CD68 (ED1) | BIO-RAD | MCA341R | mouse | mono | 1:500 |
